# S-IGTD: supervised tabular-to-image topology learning via between-group correlation for multiclass classification of biological data

**DOI:** 10.64898/2026.05.19.726105

**Authors:** Han-Ming Wu

**Affiliations:** Department of Statistics, National Chengchi University, Taipei City 11605, Taiwan, R.O.C.

**Keywords:** Convolutional neural networks, Gene expression, Feature topology optimization, Within and between analysis

## Abstract

**Motivation:** Tabular-to-image methods allow convolutional neural network (CNN)-based classifiers to analyse high-dimensional biological tables by mapping features onto a two-dimensional grid. Existing layouts are usually driven by unsupervised global correlation, which can place class-discriminative features far apart when nuisance or housekeeping covariation dominates the total covariance structure.

**Results:** We present the Supervised Image Generator for Tabular Data (S-IGTD), a supervised extension of IGTD that optimizes tabular-to-image topology by replacing total-correlation distance with one minus the absolute between-group correlation, computed from class-wise feature means, under the Within-And-Between-Analysis (WABA) decomposition. We prove entrywise consistency of the supervised distance matrix under standard moment conditions and identify balanced-class settings in which S-IGTD improves a Signal Dispersion Score (SDS)-related topology objective. In controlled simulations targeting between-group signal, S-IGTD outperformed Euclidean- and correlation-distance IGTD variants in SDS, accuracy and macro-F1 score. Across five biological benchmarks ranging from 4- to 91-class classification, S-IGTD produced compact class-supervised layouts, with 24*/*35 Holm-adjusted significant SDS wins against seven non-reference layout controls. As a secondary downstream diagnostic, a CNN with batch normalization showed higher mean accuracy than random layouts and correlation-distance IGTD on all real datasets, and higher mean accuracy than Euclidean-distance IGTD on four of five datasets, with the clearest gains on large multiclass cancer and methylation benchmarks.

**Availability and implementation:** Source code, datasets, configuration files and reproducibility scripts are freely available at https://github.com/hanmingwu1103/S-IGTD.

## 1. Introduction

High-dimensional tabular data, such as gene expression, proteomics, metabolomics and single-cell summaries, remain central to genomics and precision medicine. However, generic deep learning models do not automatically exploit the structure of such data, and strong tabular baselines such as gradient-boosted decision trees remain highly competitive (Chen and Guestrin, 2016; Ke et al., 2017; Shwartz-Ziv and Armon, 2022). For convolutional neural networks (CNNs), the central difficulty is more specific: biological feature tables have no intrinsic spatial ordering. The columns of a tabular matrix are permutation-invariant, so adjacent columns do not necessarily represent biologically or statistically neighboring features. Tabular-to-image transformations address this limitation by mapping each sample’s feature vector to a two-dimensional grid, thereby giving CNN filters a constructed local neighborhood structure on which to operate. DeepInsight (Sharma et al., 2019), REFINED (Bazgir et al., 2020) and the Image Generator for Tabular Data (IGTD) (Zhu et al., 2021) are prominent examples.

Let **X** = (*x*_*ij*_) ∈ ℝ^*n×p*^ denote a biological data matrix with *n* samples and *p* features, and let **y** ∈ {1, …, *K*}^*n*^ denote the associated class labels. A key limitation of many existing tabular-to-image layouts is that they are unsupervised: they arrange features according to global similarity in **X**, without using **y** when constructing the image topology. This can be a poor match for multiclass biological classification, where the relevant signal often lies in coordinated differences among disease, tissue or cell-type groups rather than in the strongest marginal correlations across all samples. In transcriptomic and other omics data, dominant correlations may reflect housekeeping processes (Eisenberg and Levanon, 2013), broad metabolic variation or batch-related effects, whereas the predictive signal may depend on how feature means co-vary between classes. A layout optimized for total correlation can therefore place non-discriminative background structure into local image neighborhoods while scattering class-specific covariation across the grid, where local CNN filters are less able to integrate it.

We propose the Supervised Image Generator for Tabular Data (S-IGTD), a supervised extension of IGTD that constructs feature topology from between-class rather than total covariation. The motivation is natural in biological applications because one minus the absolute correlation is already widely used as a feature dissimilarity measure, for example in co-expression analysis, but this total-correlation distance is unsupervised. S-IGTD keeps the correlation-based topology-building principle and replaces total correlation with the between-group correlation from the Within-And-Between-Analysis (WABA) decomposition (Dansereau et al., 1984). The resulting distance, one minus the absolute between-group correlation, places two features close together when their class-centroid profiles co-vary across classes. This introduces supervision at the pairwise distance-metric level and differs from supervised feature-ranking followed by row-major placement, which can prioritise important features individually but does not preserve pair-level class-related covariation.

Our contributions are methodological, theoretical and empirical. Methodologically, to the best of our knowledge, S-IGTD is the first tabular-to-image method that integrates class supervision at the pairwise distance-metric level through WABA. Theoretically, we establish consistency of the supervised distance estimator under standard moment conditions and characterise balanced-class settings in which the supervised distance improves the un-weighted local-patch surrogate tracked by the Signal Dispersion Score (SDS). Empirically, we evaluate S-IGTD in controlled simulations and on five biological benchmarks. The primary empirical endpoint is class-supervised topology compactness, measured by SDS; CNN accuracy is reported as a downstream compatibility diagnostic because the theoretical guarantee concerns feature layout rather than classifier optimality. We emphasize multiclass biological data, where class-centroid covariation is well defined and informative, while the very-small-sample dataset is included as a sensitivity case. Projection-based tabular-to-image methods are retained as important comparators because they represent a major alternative family of feature-layout generators.

The remainder of this article is organized as follows. Section 2 reviews existing tabular-to-image methods. Section 3 details the S-IGTD algorithm and the WABA-based distance metric. Section 4 derives the theoretical properties of the estimator. Sections 5 and 6 present simulation studies and real-world benchmark results, respectively, followed by conclusions in Section 7.

## 2 Background

### 2.1 Related work on tabular-to-image transformation

Existing tabular-to-image generators can be grouped into two broad families. The first family consists of projection-based methods, which embed features into a two-dimensional plane and then assign each feature to a pixel location. Sharma et al. (2019) proposed **DeepInsight**, which uses t-distributed stochastic neighbor embedding (t-SNE) or kernel principal component analysis (kernel PCA) to project features. **REFINED** (Bazgir et al., 2020) uses Bayesian multidimensional scaling (MDS), and the **TINTO** toolkit (Castillo-Cara et al., 2023; Liu et al., 2025) provides a common implementation interface for several projection-based generators. These methods are flexible, but feature collisions can occur when many features are embedded onto a small grid (Jiang et al., 2025).

The second family consists of optimization-based topology methods, represented by the Image Generator for Tabular Data (IGTD) (Zhu et al., 2021). Rather than first projecting features into a continuous two-dimensional plane, IGTD directly searches for a feature-to-pixel assignment that aligns the rank ordering of pairwise feature distances with the rank ordering of pairwise pixel distances. This rank-matching formulation avoids projection overlap, but the original IGTD distance is computed from the full data matrix without using class labels. Thus, both projection-based generators and standard IGTD usually construct image topology in an unsupervised manner.

Two related literatures clarify what kind of supervision is missing. First, supervised *feature-ranking* methods, including linear discriminant rank (Li et al., 2004), minimum-redundancy-maximum-relevance (mRMR) (Peng et al., 2005), ReliefF (Robnik-Šikonja and Kononenko, 2003) and Boruta wrappers (Kursa and Rudnicki, 2010), order features by class-discriminative importance. If such rankings are combined with row-major placement, labels enter the construction only through univariate feature scores; pairwise class-related covariation is not preserved. Second, non-grid tabular deep-learning architectures, including Tab-Net (Arik and Pfister, 2021), FT-Transformer (Gorishniy et al., 2021), graph-based tabular models and prior-data fitted networks such as TabPFN (Hollmann et al., 2023), avoid the image-layout problem entirely. These approaches are complementary to tabular-to-image methods, which remain useful when a two-dimensional feature topology is desired for CNN filters, interpretable local neighborhoods or compatibility with vision-based architectures. S-IGTD differs from these approaches by introducing class supervision directly into the pair-wise distance metric used to construct the image topology.

### 2.2 Problem formulation and IGTD

Using the notation introduced above, let **X** = (*x*_*ij*_) ∈ ℝ^*n×p*^ denote a data matrix with *n* samples and *p* features, and let **y** ∈ {1, …, *K*}^*n*^ denote the associated class labels. We write **x**_*i*_ = (*x*_*i*1_, …, *x*_*ip*_)^⊤^ for the feature vector of sample *i* and **X**_·*j*_ = (*x*_1*j*_, …, *x*_*nj*_)^⊤^ for feature column *j*. The objective is to map each sample vector **x**_*i*_ to an image 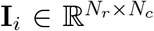, where *N*_*r*_*N*_*c*_ = *p*.

The core challenge is to find a permutation

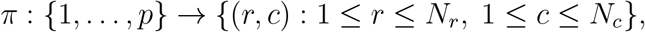

which assigns each feature to a unique pixel coordinate. We distinguish two layout paradigms. In an *unsupervised* arrangement, *π* is optimized from a feature distance

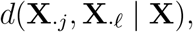

which depends only on the marginal feature structure in **X**. In a *supervised* arrangement, *π* is optimized from a conditional feature distance

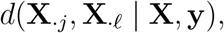

which also depends on the class labels. Existing tabular-to-image methods such as DeepInsight, REFINED and IGTD primarily operate under the unsupervised paradigm. Supervised pre-processing variants may use labels before the layout step, usually through univariate feature ranking, but the spatial layout itself is not constructed from pairwise class-related covariation.

The Image Generator for Tabular Data (IGTD) (Zhu et al., 2021) instantiates the unsupervised paradigm as a Quadratic Assignment Problem (QAP). It assigns features to a fixed pixel grid by aligning the rank ordering of pairwise feature distances with the rank ordering of pairwise pixel distances. This creates local image neighborhoods in which nearby pixels tend to contain features that are close under the chosen feature distance. The algorithm constructs two *p* × *p* rank matrices:

1. **Feature distance rank matrix (R):** This matrix is derived from pairwise distances between feature columns. If *d*(**X**_·*j*_, **X**_·*ℓ*_) is the distance between features *j* and *ℓ*, then **R**_*jℓ*_ is the rank of this distance among all feature pairs.
2. **Pixel distance rank matrix (Q):** This matrix is derived from spatial Euclidean distances between pixel coordinates on the target grid. If *π*(*j*) and *π*(*ℓ*) are the pixel coordinates assigned to features *j* and *ℓ*, then **Q**_*π*(*j*),*π*(*ℓ*)_ is the rank of their physical grid distance among all pixel pairs.

IGTD seeks a permutation *π* that aligns these two rank structures by minimizing

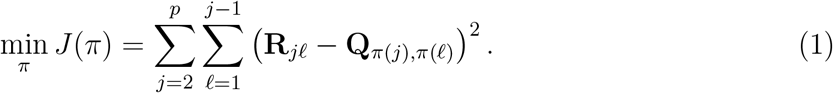

The original IGTD constructs **R** using either Euclidean distance (IGTD-ED) or a correlation-based distance (IGTD-RD) computed over the entire data matrix **X**, so the class labels **y** do not affect the topology. The optimized permutation *π* is then applied to every sample to produce the image **I**_*i*_. S-IGTD modifies this construction at exactly one point: the feature distance used to build **R**.

## 3 Method

In this section, we introduce the Supervised Image Generator for Tabular Data (S-IGTD). The method is motivated by the Within-And-Between-Analysis (WABA) decomposition and implemented as a supervised replacement of the feature-distance matrix used by IGTD. We first describe the WABA-based sample correlation decomposition, then define the S-IGTD distance, algorithm and practical considerations for high-dimensional biological data.

### 3.1 The supervised IGTD (S-IGTD)

S-IGTD makes a targeted but statistically substantive modification to IGTD: it replaces the unsupervised pairwise feature distance used to construct the rank matrix **R** with a supervised distance based on between-class covariation. Standard IGTD computes feature distances from Euclidean distance or total correlation in the raw data matrix **X**. In supervised classification, however, total correlation combines between-class structure with within-class covariation, and the latter may reflect nuisance variation rather than discriminative signal.

This construction is motivated by common practice in biological data analysis, where one minus the absolute correlation is widely used as a feature dissimilarity measure, for example in co-expression analysis (Langfelder and Horvath, 2008). S-IGTD preserves this correlation-based topology-building principle but replaces total correlation with the between-class correlation from the WABA decomposition (Dansereau et al., 1984). Thus, feature pairs are placed close together when their class-centroid profiles co-vary across classes.

Let **X** = (*x*_*ij*_) ∈ ℝ^*n×p*^ denote the training data matrix, and let **y** = (*y*_1_, …, *y*_*n*_)^⊤^ ∈ {1, …, *K*}^*n*^ denote the class labels. For feature indices *j, ℓ* ∈ {1, …, *p*}, let **X**_·*j*_ and **X**_·*ℓ*_ denote the corresponding columns of **X**. We write 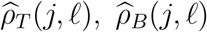 and 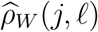 for the empirical total, between-class and within-class correlations associated with the observed feature pair (**X**_·*j*_, **X**_·*ℓ*_).

For a discrete class variable, WABA decomposes the sample total correlation between two feature columns into a between-class component and a within-class component. Following the WABA notation, the decomposition can be written as

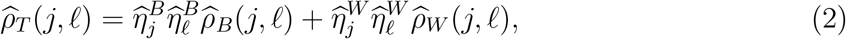

where 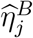 and 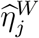 are the sample between- and within-eta coefficients for feature column **X**._*j* ._The empirical between-class correlation 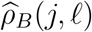 is the correlation of the class-level means of feature columns **X**_·*j*_ and **X**_·*ℓ*_, whereas 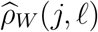 is the correlation of within-class deviations after removing class-level effects.

The WABA decomposition motivates the S-IGTD distance. A topology based on 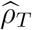 may co-localize features whose association is driven by within-class housekeeping, batch or background variation. S-IGTD instead uses the between-class component, through class-balanced centroid correlation, to prioritise feature pairs that co-vary across class means. The conditions under which this improves the topology objective are given in Section 4.

For implementation, S-IGTD first constructs a class-centroid matrix

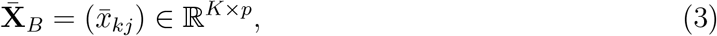

where 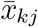 is the sample mean of feature column **X**_·*j*_ in class *k*. Let *w*_1_, …, *w*_*K*_ be nonnegative class weights satisfying 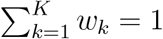, and let

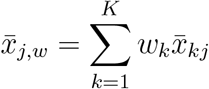

be the weighted centroid mean for feature column **X**_·*j*_. The weighted empirical between-class correlation, 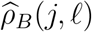, used by S-IGTD is

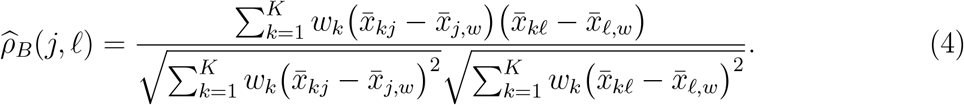

The choice *w*_*k*_ = *n*_*k*_*/n*, where *n*_*k*_ is the number of samples in class *k*, gives the usual sample-size-weighted WABA between-class correlation. The choice *w*_*k*_ = 1*/K* gives the class-balanced version used by S-IGTD in the main analyses. Unless otherwise stated, 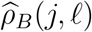 refers to this class-balanced version.

The empirical S-IGTD supervised distance is

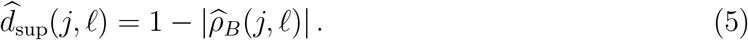

The absolute value allows both positively and negatively coordinated class-centroid profiles to be treated as close in the image topology. This is useful for CNN filters, which can learn signed local contrasts, but the formal SDS guarantee in Section 4 applies directly to positive between-class correlations. The use of the absolute value is therefore a modeling choice rather than a direct consequence of the SDS proof. The complete procedure is summarized in Algorithm 1.

#### Algorithm 1 Supervised Image Generator for Tabular Data (S-IGTD)

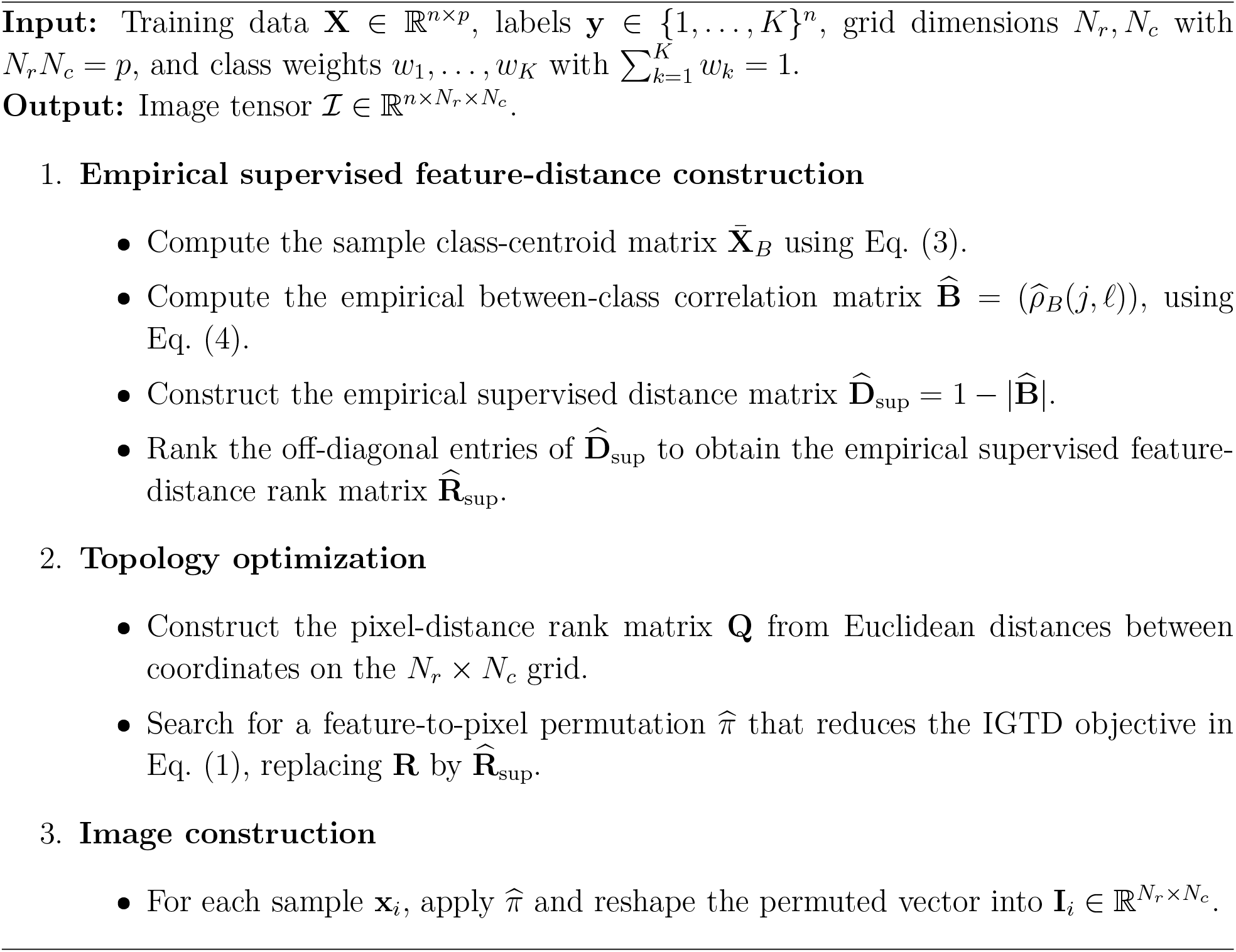

### 3.2 Practical considerations

S-IGTD is designed for high-dimensional biological data, where the number of features is often large, class sizes may be unbalanced and some features may show little or no between-class variation.

#### 3.2.1 Computational complexity

For correlation-distance IGTD, the feature-distance step scales as *O*(*np*^2^) because correlations are computed from all *n* samples. S-IGTD replaces this step with correlation computation on the *K* × *p* sample centroid matrix 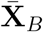, which scales as *O*(*Kp*^2^). Since *K* is usually much smaller than *n* in biological classification tasks, the supervised distance construction is not the computational bottleneck. The dominant cost remains the QAP search step, which is shared by IGTD and S-IGTD.

#### 3.2.2 Class imbalance and centroid stabilization

Class imbalance is common in biomedical datasets. The sample-size-weighted WABA version uses *w*_*k*_ = *n*_*k*_*/n*, so large classes contribute more strongly to the between-class correlation. In contrast, the default S-IGTD construction uses *w*_*k*_ = 1*/K*, so each class contributes equally to the feature topology. This class-balanced choice is useful when minority classes are scientifically important and should not be dominated by majority classes during layout construction.

For small or noisy classes, a single class centroid may be unstable. As an optional stabilization step, S-IGTD can augment the centroid matrix by repeatedly subsampling observations within each class and computing additional within-class centroids. These bootstrap centroids increase the number of rows used to estimate between-class correlation and can reduce sensitivity to individual observations in small classes. The exact estimator analyzed in Section 4 is the basic *K* × *p* centroid estimator; the bootstrap-stabilized version is an empirical robustness variant used for practical deployment.

#### 3.2.3 Features with negligible between-class variation

Some biological features have zero or near-zero variation across class centroids. For such features, 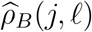 is undefined or unstable because the denominator in Eq. (4) is zero or close to zero. In the default rule, feature pairs involving a zero-between-variation feature are assigned the maximum supervised distance, 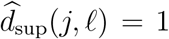. This discourages such features from being placed close to discriminative feature groups.

A softer alternative is to shrink the empirical supervised similarity by the amount of estimated between-class variation. Define

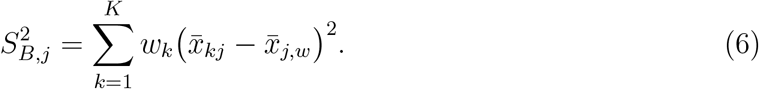

Then the shrinkage version of the distance is

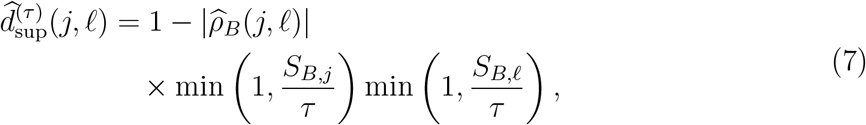

where *τ >* 0 is a small threshold. This rule downweights empirical correlations involving features with negligible between-class variation. The main empirical benchmarks use the hard zero-variation rule, while Eq. (7) provides a sensitivity option.

#### 3.2.4 Binary classification

The exact S-IGTD estimator is most informative when *K* ≥ 3. When *K* = 2, each feature with nonzero between-class variation has a two-point centroid vector

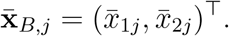

For any pair *j, ℓ* with 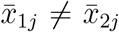 and 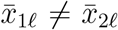,the Pearson correlation of two nonconstant two-vectors satisfies

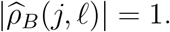

Consequently,

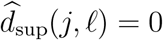

for every such pair. Pairs involving a feature with zero between-class variation are assigned 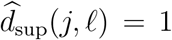 by the singularity rule above. Thus, the supervised distance collapses to a two-level matrix, and the QAP solver loses the continuous ranking information needed for fine within-signal ordering. This is a geometric property of two-point Pearson correlation, not a degrees-of-freedom argument. The exact theoretical results below therefore focus on multiclass settings with *K* ≥ 3.

## 4 Theoretical properties

In this section, we analyse the statistical properties of the exact multiclass S-IGTD estimator. Section 3 defined the empirical construction in terms of observed feature columns **X**_·*j*_ and **X**_·*ℓ*_. Here, *X*_*j*_ and *X*_*ℓ*_ denote the corresponding population feature variables. We use *ρ*_*T*_ (*j, ℓ*), *ρ*_*B*_(*j, ℓ*) and *ρ*_*W*_ (*j, ℓ*) for the population total, between-class and within-class correlations associated with the feature pair (*X*_*j*_, *X*_*ℓ*_), and we use hatted quantities such as 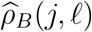 and 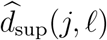 for their empirical counterparts.

Throughout this section, we assume that (i) *K* ≥ 3, with the binary degeneracy described in Section 3.2; (ii) the between-class variance of every feature involved in a statement is strictly positive unless the singularity convention in Section 3.2 is invoked; and (iii) the class-balanced centroid correlation used by S-IGTD is the population counterpart of Eq. (4) with *w*_*k*_ = 1*/K*. When classes are balanced, this class-balanced correlation coincides with the probability-weighted WABA between-class correlation; when classes are unbalanced, it is a class-balanced population surrogate.

We write

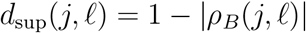

for the population supervised distance and

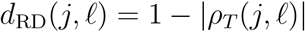

for the population analogue of the correlation-distance IGTD baseline. The use of the absolute value has two motivations. First, it treats positively and negatively coordinated class-centroid profiles as equally local in the topology, which is useful when a local image filter can learn signed contrasts between neighboring features. Second, it keeps the supervised distance on the same [0, 1] scale as the usual absolute-correlation distance used in correlation-based feature layouts. The formal unweighted-sum patch argument in Section 4.3 directly supports positive between-class correlations; negatively correlated local adjacencies are justified as a modeling choice for contrast-based local aggregation rather than as a consequence of that proposition alone.

The results below establish entrywise consistency of the supervised distance matrix and identify biologically interpretable scenarios in which the supervised metric gives a more appropriate local adjacency priority than total-correlation distance. We do not claim consistency of the downstream QAP permutation: that would require additional identifiability and optimization assumptions beyond the distance-estimation problem considered here. The relationship between these topology results and classifier performance is evaluated empirically in both the simulation studies and the real-data analysis.

### 4.1 Asymptotic consistency

A fundamental requirement for a statistical topology generator is that its estimated feature-distance matrix converges entrywise to the intended population distance. The following result establishes this property for the S-IGTD distance.

#### Theorem 1 (Consistency of the S-IGTD distance).

*Let* 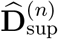 *be the empirical supervised distance matrix computed from a dataset of size n with K classes, using the class-balanced weights w*_*k*_ = 1*/K in Eq*. (4). *Fix a feature pair* (*j, ℓ*) *such that*

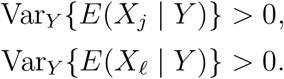

*If K is fixed and n*_*k*_ → ∞ *for every class k, then*

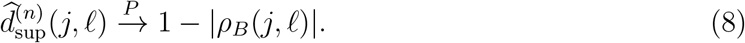

*If* Var_*Y*_ {*E*(*X*_*j*_ | *Y* )} = 0 *or* Var_*Y*_ {*E*(*X*_*ℓ*_ | *Y* )} = 0, *then under the singularity convention in Section 3.2*,

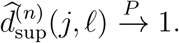

*Proof*. Let

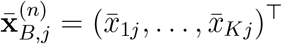

denote the vector of sample class means for feature *j*, and let

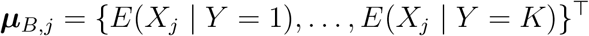

denote its population counterpart. By the Weak Law of Large Numbers, as *n*_*k*_ → ∞ for every class *k*,

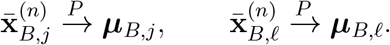

On the set where both population between-class variances are non-zero, the weighted Pearson correlation map with fixed weights *w*_*k*_ = 1*/K* is continuous. Therefore, by the Continuous Mapping Theorem,

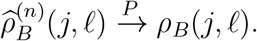

Applying the continuous map *x* ↦ 1 − |*x*| gives Eq. (8). If either feature has zero between-class variance, the centroid correlation is singular, and the rule in Section 3.2 assigns 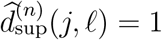, giving the stated limit.

### 4.2 WABA decomposition and topological divergence

We next compare the adjacency priorities induced by *d*_RD_ and *d*_sup_. The relevant population analogue of the WABA decomposition in Eq. (2) is

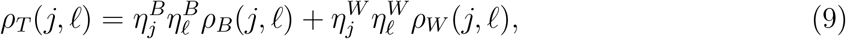

where 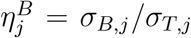 and 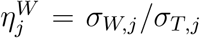 are the between- and within-eta coefficients for feature *j*. The following scenarios are stated in the balanced-class population setting, where the class-balanced S-IGTD correlation coincides with the standard WABA between-class correlation. They illustrate when the supervised and unsupervised distances induce different local adjacency priorities in the QAP objective.

The practical interpretation is important in biological data. In omics classification, a large total correlation *ρ*_*T*_ may arise from at least two different sources: coordinated class-level shifts that are useful for distinguishing disease, tissue or cell-type groups, or within-class covariation caused by housekeeping processes, metabolic background, batch effects or other nuisance structure. S-IGTD is designed for the first source. The four scenarios below translate this decomposition into feature-layout consequences.

#### 4.2.1 Scenario 1: signal attenuation, the “damping” effect

Consider a pair of discriminative features whose between-class covariation is strong but whose total correlation is attenuated by within-class variation. Biologically, this may occur for co-regulated biomarkers whose class-centroid profiles move together across disease subtypes or tissue groups, but whose sample-level variation is noisy within each class. For example, two genes may show coordinated up- or down-regulation across cancer types, yet their total correlation can be damped by patient heterogeneity, measurement noise or within-tissue regulatory variation.

##### Condition 1

(Signal attenuation).

Let |*ρ*_*B*_(*j, ℓ*)| = 1, *ρ*_*W*_ (*j, ℓ*) = 0, and 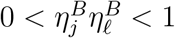.

##### Corollary 2

(Adjacency recovery under signal attenuation). *Under Condition 1*,

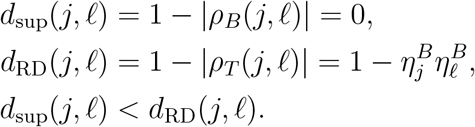

This follows immediately from Eq. (9). The supervised distance gives this discriminative pair the highest adjacency priority, whereas the total-correlation distance assigns it a strictly positive distance. In layout terms, S-IGTD is less likely to separate class-informative feature pairs simply because their total sample-level correlation has been weakened by within-class variability.

#### 4.2.2 Scenario 2: noise dominance, the “housekeeping” effect

Consider features that covary strongly within classes but show little or no between-class covariation. This scenario is common in biological data. Housekeeping genes, broad metabolic programs, ribosomal or mitochondrial activity, sample quality effects and batch-related processes can create strong correlations across all samples or within biological groups without carrying useful class-discriminative information. A total-correlation layout may incorrectly place such features close together because they are strongly correlated, even though their association is not relevant to the supervised classification task.

##### Condition 2

(Noise dominance).

Let |*ρ*_*W*_ (*j, ℓ*)| = 1 and suppose 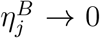 and 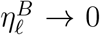. If *ρ*_*B*_(*j, ℓ*) is defined, assume *ρ*_*B*_(*j, ℓ*) = 0; if the between-class variance of either feature is zero, adopt the singularity convention *d*_sup_(*j, ℓ*) = 1 from Section 3.2.

##### Corollary 3

(False adjacency under noise dominance). *Under Condition 2*,

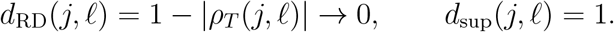

Substituting Condition 2 into Eq. (9) gives |*ρ*_*T*_ (*j, ℓ*)| → 1. The total-correlation distance therefore gives this pair high adjacency priority, whereas the supervised distance assigns it maximum feature distance and discourages local adjacency driven purely by within-class covariation. This is the main bioinformatics motivation for replacing total correlation with between-class correlation: the layout should not spend local CNN neighborhood capacity on background biological or technical structure that does not separate the classes.

#### 4.2.3 Scenario 3: Simpson’s paradox, signal cancellation

A third failure mode occurs when between-class and within-class associations have opposite signs and cancel in the total correlation. In biological applications, this can arise when disease- or tissue-level shifts induce one association pattern across class centroids, while within-class regulation induces the opposite association among individual samples. For example, two features may increase together across tumour types, but within each tumour type one feature may compensate for the other, producing a negative within-class association. The total correlation can then be close to zero, even though the class-centroid association is strong.

##### Condition 3 (Simpson’s paradox).

Let *ρ*_*B*_(*j, ℓ*) = 1 and *ρ*_*W*_ (*j, ℓ*) = −1.

If, in addition,

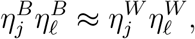

then Eq. (9) gives *ρ*_*T*_ (*j, ℓ*) ≈ 0. Hence

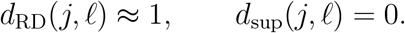

The supervised distance recovers the between-class adjacency priority that total correlation masks under Condition 3. This scenario is especially relevant when class structure is the scientific object of interest: total correlation may fail to detect class-level coordination because opposing within-class effects cancel it.

#### 4.2.4 Scenario 4: equivalence, between-class dominance

The supervised and total-correlation distances coincide when between-class variation dominates the total variation. This is the favorable case for unsupervised correlation layouts. Biologically, it corresponds to features whose dominant source of variation is the class label itself, such as strong marker genes, methylation loci or proteomic features that are nearly constant within a class but differ systematically between classes.

##### Condition 4 (Equivalence).

Let 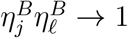 and 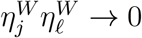.

Under Condition 4, Eq. (9) gives

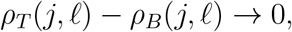

and therefore

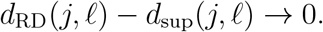

Thus, when nearly all relevant variation is between classes, total-correlation IGTD and S-IGTD induce the same pairwise distance ordering for that feature pair. This scenario clarifies that S-IGTD is not expected to differ from correlation-distance IGTD when the global correlation structure is already dominated by supervised class separation.

Table 1 summarizes the four scenarios.

**Table 1.**
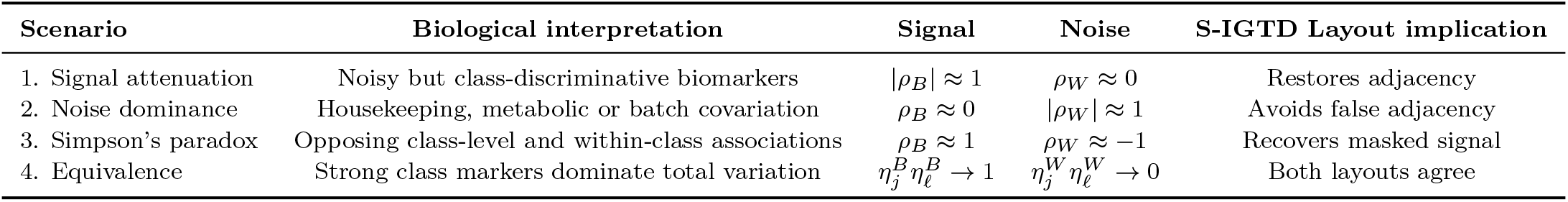
Comparison of topological distances under four WABA scenarios. Here *d*_RD_ = 1 − |*ρ*_*T*_ | is the total-correlation distance and *d*_sup_ = 1 − |*ρ*_*B*_| is the S-IGTD supervised distance.

### 4.3 Maximisation of unweighted-sum patch discriminability

We now connect the between-class distance to a simple local aggregation objective. The Signal Dispersion Score (SDS), formally defined in Section 5, measures how spatially compact a set of signal features is under a learned feature-to-pixel permutation. The result below does not prove optimality of SDS itself. Instead, it shows why placing positively between-correlated signal features in the same local neighborhood can increase a simple between-class patch signal.

Let Ω denote the set of feature indices in a local image neighborhood, for example a 3 × 3 patch. Define the aggregate patch signal as the unweighted sum

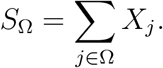

To quantify class separation in this patch readout, define

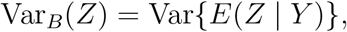

and let *σ*_*B,j*_ = {Var_*B*_(*X*_*j*_)}^1*/*2^ denote the between-class standard deviation of feature *j*.

#### Proposition 1 (Unweighted-sum patch discriminability).

For a fixed patch membership Ω with fixed between-class standard deviations {*σ*_*B,j*_ : *j* ∈ Ω},

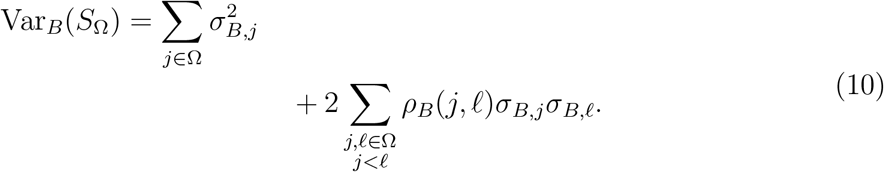

Hence, if *ρ*_*B*_(*j, ℓ*) ≥ 0 for all pairs in Ω, the between-class variance of the unweighted patch sum is increased by grouping feature pairs with large positive between-class correlations.

*Proof*. Let *Z*_*j*_ = *E*(*X*_*j*_ | *Y* ). Then

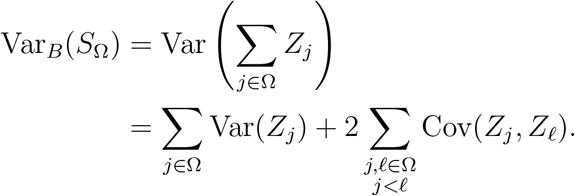

By definition,

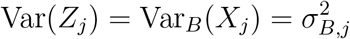

and

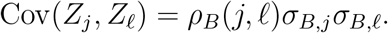

Substitution gives Eq. (10).

Equation (10) directly supports the S-IGTD distance for positively correlated between-class signal pairs. The use of |*ρ*_*B*_| extends this principle to negatively coordinated class-centroid profiles. This extension is not implied by the unweighted-sum objective, because negative correlations reduce the variance of an unweighted sum. Its motivation is instead contrast-based: local image filters, especially CNN filters, can assign opposite signs to neighboring pixels and therefore exploit strong negative as well as strong positive class-level coordination. Thus, 1 − |*ρ*_*B*_| should be interpreted as a topology-building distance for local contrast learning, while Proposition 1 provides a formal guarantee for the positive-correlation case.

The proposition also clarifies the scope of the theory. The unweighted sum is not the feature map learned by a general two-dimensional image classifier, nor by a CNN kernel, which applies learned weights and nonlinearities. Therefore, the theory supports the topology objective rather than classifier optimality. This is why SDS is treated as the primary topology metric in Sections 5 and 6, whereas accuracy and the macro-F1 score are reported as downstream diagnostics.

## 5 Simulation studies

We conducted four controlled simulation studies to evaluate S-IGTD under settings that isolate the supervised replacement of the IGTD feature-distance metric. Study 1 evaluates robustness to nuisance correlation and is presented first because it targets the main biological failure mode motivating the method: label-irrelevant nuisance or “housekeeping” correlation dominating an unsupervised topology. Study 2 examines finite-sample behavior related to Theorem 1. Study 3 uses label permutation to test whether the learned topology depends on the class labels. Study 4 stresses joint growth in feature dimension and signal mass. The Simpson’s-paradox scenario in Section 4 is treated analytically rather than as a separate Monte Carlo study, because it requires a deliberately engineered sign cancellation between between-class and within-class correlations.

The comparator set is deliberately restricted to IGTD-ED, IGTD-RD and S-IGTD. This restriction is intentional: the simulation purpose is to isolate the effect of replacing IGTD’s unsupervised feature distance with the between-class WABA distance. Projection-based tabular-to-image methods are evaluated in the real-data benchmark, where comparison across tabular-to-image families is more meaningful. All simulations use the multiclass S-IGTD estimator from Section 3: class-balanced centroid correlation with *w*_*k*_ = 1*/K* and no centroid bootstrapping. Studies 1 and 3 use the default singularity rule for zero-between-variation features, whereas Studies 2 and 4 use the soft singularity rule in Eq. (7) with *τ* = 0.05.

### 5.1 Common data-generating model and evaluation

All simulation studies use *K* = 6 balanced classes. In each Monte Carlo replicate, class labels are generated with exactly equal class sizes: for *k* ∈ {1, …, 6}, class *k* contributes *n*_*k*_ samples, so *n* = 6*n*_*k*_ and *n*_1_ = · · · = *n*_6_ = *n*_*k*_. Thus the labels are fixed to be balanced rather than sampled from a multinomial distribution; in the implementation, the training labels are randomly permuted after construction before feature generation. The feature set {1, …, *p*} is divided into a ground-truth signal set 𝒮_sig_ ⊂ {1, …, *p*} and its nuisance complement. The signal set is further partitioned into three equal-size blocks,

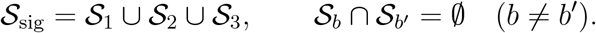

In Studies 1–3, the signal-block size is fixed by the study design. In Study 4, the signal fraction is fixed at 12% of *p*.

For a signal feature *j* ∈ 𝒮_*b*_, the observation for sample *i* is generated from a block-specific class-centroid shift plus independent noise,

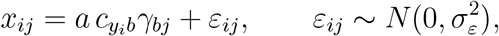

where *c*_·*b*_ is a fixed class-profile vector for block *b, a* controls the signal amplitude, 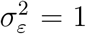, and *γ*_*bj*_ is a feature-specific signal coefficient. For Studies 1, 2 and 4,

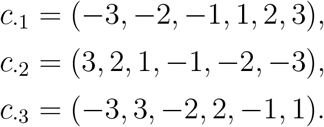

For Study 3,

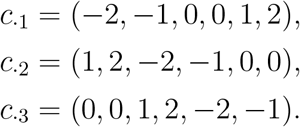

The notation 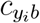 means that the class label *y*_*i*_ selects the corresponding entry of the class-profile vector for block *b*. Equivalently, for *j* ∈ 𝒮_*b*_ and *y*_*i*_ = *k*,

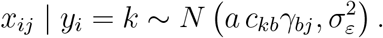

Thus the relationship between the label and a signal feature is entirely through the class-specific mean shift *a c*_*kb*_*γ*_*bj*_. Study 3 fixes *γ*_*bj*_ = 1. Studies 1, 2 and 4 draw *γ*_*bj*_ independently as a log-normal magnitude, exp{*N* (0, 0.9^2^)}, multiplied by a random sign; the sign-flip probabilities are 0.5, 0.9 and 0.9, respectively. The signal amplitudes are *a* = 1.6 in Studies 1 and 2, *a* = 1.0 in Study 3 and *a* = 1.3 in Study 4.

Nuisance features have no systematic class-centroid shift; their conditional mean is zero for every class, so they do not depend on *y*_*i*_ through the class mean. They are partitioned into exchangeable covariance blocks by the allocation rule used in the simulator: with *p*_noise_ = *p* −|𝒮_sig_| and nominal block size *q*, the number of nuisance blocks is *M* = max{1, ⌊*p*_noise_*/q*⌋}, block sizes are as equal as possible, and any remainder is assigned to the first blocks. For nuisance block *m*,

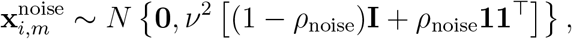

where *ν* is the nuisance scale, so each nuisance feature has marginal variance *ν*^2^ before any added latent effects. Studies 1, 2 and 4 additionally add class-independent sample latent effects to nuisance features only: 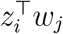 is generated from independent standard-normal factors and loadings, standardized featurewise, and multiplied by a latent scale. This construction creates strong nuisance correlation controlled by *ρ*_noise_ and latent factors while keeping nuisance features non-discriminative at the class-centroid level. The same generated training and test sets are used for IGTD-ED, IGTD-RD and S-IGTD within each Monte Carlo replicate. The random-state seeds are 42 for Studies 1 and 4 and 10 for Studies 2 and 3, with replicate seeds obtained by deterministic setting-specific offsets.

All three layout methods produce a feature-to-pixel permutation. The resulting simulation images are evaluated with a lightweight LeNet-5 CNN diagnostic trained from scratch. This differs from the Conv-BN-ReLU classifier used for the real-data benchmark in Section 6. Classifier outcomes are therefore secondary diagnostics in the simulations; the primary simulation endpoint is topology compactness measured by SDS.

We report three outcomes:

- **Signal Dispersion Score (SDS):** topology compactness of true signal features; lower is better.
- **Accuracy:** test-set classification accuracy.
- **Macro-F1 score:** the unweighted mean of class-specific F1 scores; higher is better.

For a learned feature-to-pixel permutation 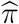, let

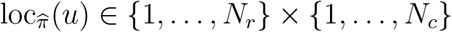

denote the grid coordinate assigned to feature *u*. The Signal Dispersion Score is the average pairwise Euclidean distance among the true signal features:

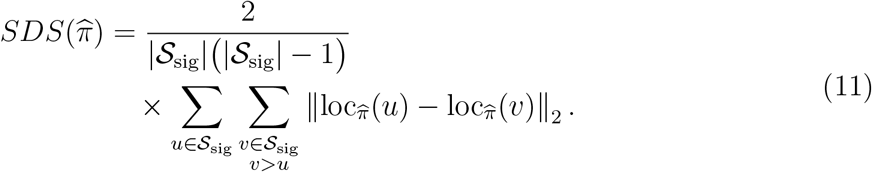

On simulated data, 𝒮_sig_ is known from the generator. On real data, where ground-truth signal features are unavailable, it is replaced by the ANOVA-*F* proxy described in Section 6.

For each simulation setting and outcome metric, paired Wilcoxon signed-rank tests compare S-IGTD with IGTD-ED and IGTD-RD across matched Monte Carlo replications. One-sided alternatives are used, with “less” for SDS and “greater” for accuracy and macro-F1 score. Within each setting and metric, the two *p*-values are adjusted by the Holm-Bonferroni procedure (Holm, 1979). We use “Holm-adjusted significant” to mean Holm-adjusted *p <* 0.05.

Figures 1 and 2 provide qualitative snapshots of the generated image representations and signal-feature locations. Figures 3–5 provide the quantitative trends across simulation settings, and Table 2 summarizes the replicate means.

**Table 2.** Simulation summary over Monte Carlo replications. Values are means over matched replications for each setting. Lower SDS and higher accuracy favor a method. Paired tests are described in the text and are reproducible from the per-replication result tables.

| Study | Setting | Reps | SDS |  |  | Accuracy |  |  |
| --- | --- | --- | --- | --- | --- | --- | --- | --- |
|  |  |  | IGTD-ED | IGTD-RD | S-IGTD | IGTD-ED | IGTD-RD | S-IGTD |
| 1 | $\rho_{\text{noise}} = 0.80$ | 30 | 8.33 | 10.57 | 7.62 | 0.957 | 0.969 | 0.978 |
| 1 | $\rho_{\text{noise}} = 0.85$ | 30 | 8.37 | 10.87 | 7.63 | 0.959 | 0.973 | 0.980 |
| 1 | $\rho_{\text{noise}} = 0.90$ | 30 | 8.31 | 10.69 | 7.70 | 0.957 | 0.969 | 0.983 |
| 1 | $\rho_{\text{noise}} = 0.95$ | 30 | 8.37 | 10.57 | 7.76 | 0.962 | 0.972 | 0.981 |
| 2 | $n_k = 20$ | 30 | 14.17 | 16.01 | 11.28 | 0.894 | 0.910 | 0.946 |
| 2 | $n_k = 50$ | 30 | 14.33 | 15.86 | 11.25 | 0.950 | 0.966 | 0.980 |
| 2 | $n_k = 100$ | 30 | 13.91 | 15.53 | 11.33 | 0.976 | 0.988 | 0.996 |
| 2 | $n_k = 200$ | 30 | 13.45 | 15.52 | 11.25 | 0.985 | 0.993 | 0.998 |
| 4 | $p = 900$ | 10 | 13.27 | 15.79 | 11.36 | 0.790 | 0.822 | 0.877 |
| 4 | $p = 1600$ | 10 | 19.40 | 20.07 | 14.39 | 0.900 | 0.935 | 0.948 |
| 4 | $p = 2500$ | 10 | 24.26 | 25.10 | 18.39 | 0.934 | 0.944 | 0.970 |

**Figure 1.**
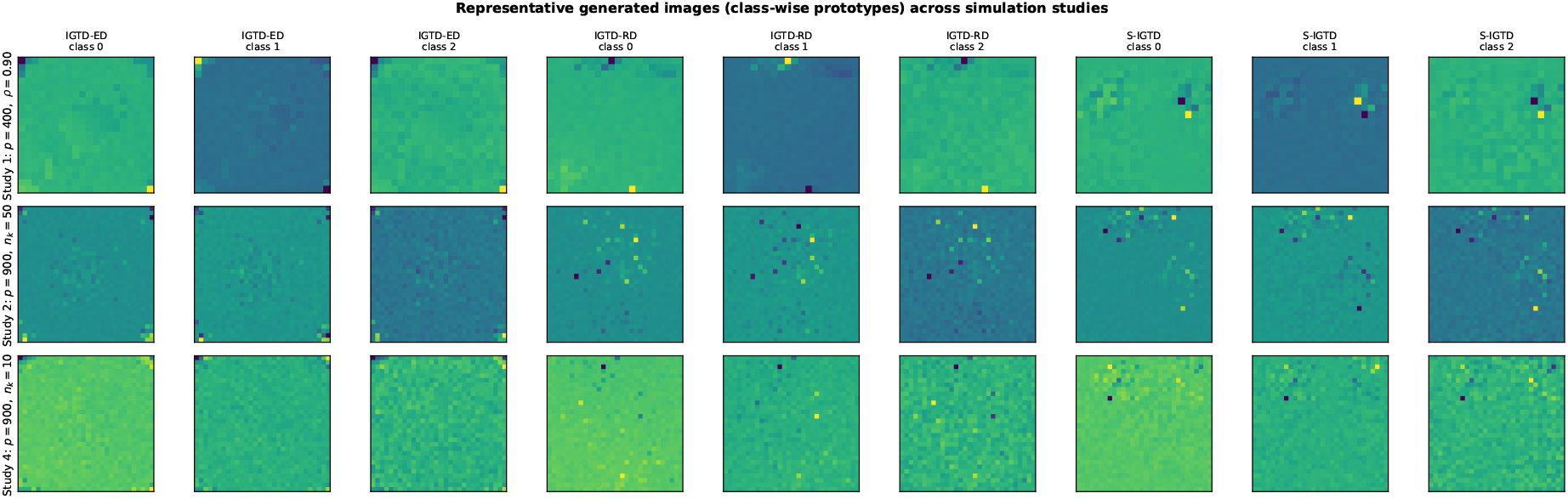
Representative generated images, shown as class-wise prototypes across simulation studies. Rows correspond to representative settings from Studies 1, 2 and 4; columns compare IGTD-ED, IGTD-RD and S-IGTD for selected classes. Only three of the six classes are displayed for legibility. These images are qualitative examples rather than the main evidence; quantitative topology and classifier trends are reported in Figures 2–5 and Table 2.

**Figure 2.**
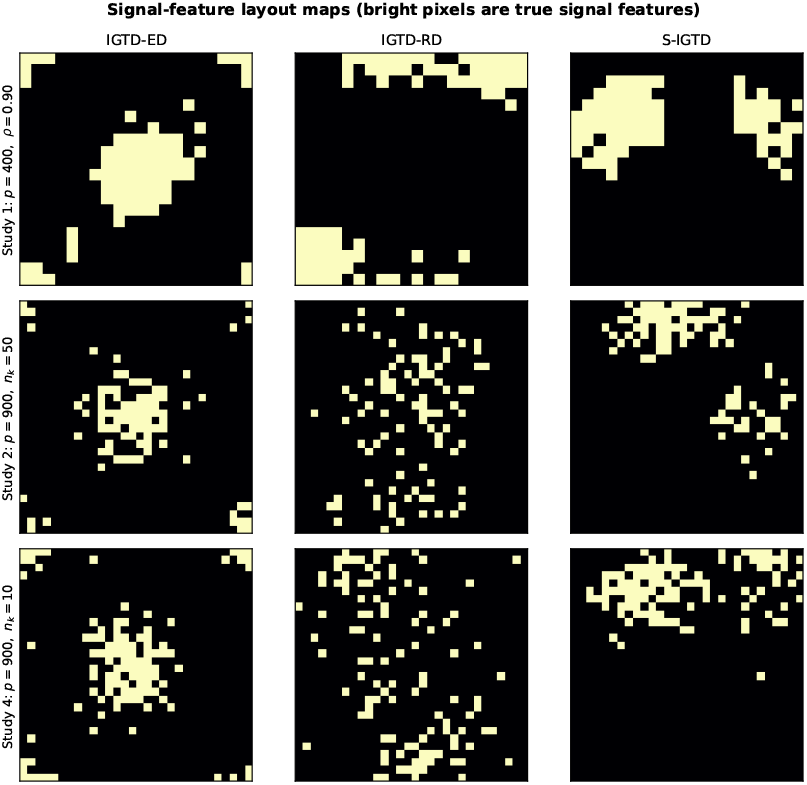
Signal-feature layout maps for the same representative Study 1/2/4 settings shown in Figure 1. Bright pixels indicate true signal features. S-IGTD may show two or more compact subregions because the simulator contains multiple signal blocks with different class-centroid profiles. SDS measures the average pairwise distance among signal-feature coordinates and does not require a single connected component.

**Figure 3.**
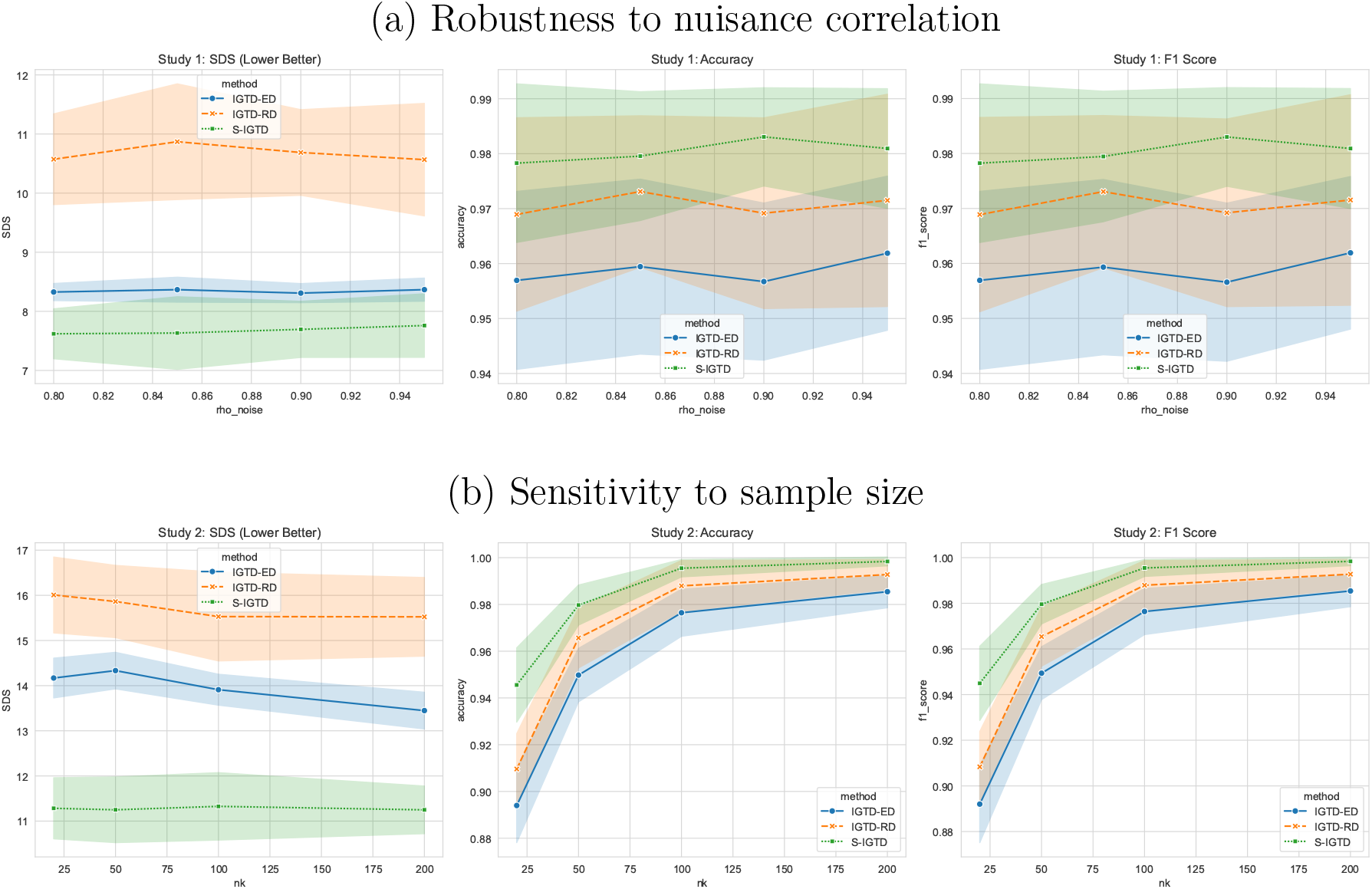
Simulation trends for Studies 1 and 2. Panel (a) varies nuisance correlation *ρ*_noise_; panel (b) varies the number of training samples per class *n*_*k*_. Lines show Monte Carlo means across replications; shaded bands are the seaborn default 95% confidence intervals around the mean. Numerical summaries are reported in Table 2.

Figure 1 shows generated class-prototype images, whereas Figure 2 removes intensity and shows only signal-feature locations for the same representative settings.

### 5.2 Study 1: robustness to nuisance correlation

#### Design

Study 1 targets the noise-dominance scenario from Section 4. We set *p* = 400 with a 20 × 20 grid, 72 signal features arranged as three blocks of 24, *n*_train_ = 180 with 30 samples per class, and *n*_test_ = 600 with 100 samples per class. Nuisance features used nominal block size 80, giving four nuisance blocks of 82 features, nuisance scale *ν* = 3.0, ten class-independent latent factors with latent scale 5.0, and the nuisance-correlation level was varied as

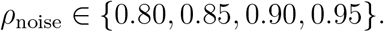

Each setting used 30 Monte Carlo replications. Increasing *ρ*_noise_ strengthens within-class nuisance correlation among non-signal features while leaving the class-centroid signal un-changed.

#### Result interpretation

Figure 3(a) shows that S-IGTD has lower mean SDS than both IGTD variants at every nuisance-correlation level. Lower SDS means that the true signal features remain spatially compact under S-IGTD even when strong within-class nuisance correlation is present. Accuracy and macro-F1 score also remain higher in mean for S-IGTD, but these classifier outcomes are secondary to the topology result. The paired comparisons are Holm-adjusted significant in the released result tables. The stable SDS gap as *ρ*_noise_ increases is consistent with the noise-dominance argument in Corollary 3.

### 5.3 Study 2: sensitivity to sample size

#### Design

Study 2 examines finite-sample behavior around the consistency result in Theorem 1. We used *p* = 900 with a 30 ×30 grid, 90 signal features arranged as three blocks of 30, and fixed *n*_test_ = 600 with 100 samples per class. Nuisance features used nominal block size 120, giving six nuisance blocks of 135 features, nuisance scale *ν* = 4.5, four class-independent latent factors with latent scale 2.5, and *ρ*_noise_ = 0. The number of training samples per class was varied as

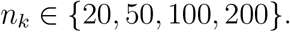

Each setting used 30 Monte Carlo replications.

#### Result interpretation

Figure 3(b) shows that increasing *n*_*k*_ improves estimation and classifier outcomes, especially for accuracy and macro-F1 score. S-IGTD nevertheless maintains the lowest mean SDS at every sample size, indicating that the between-class centroid correlation remains useful even at moderate training sizes. The paired comparisons are Holm-adjusted significant in the released result tables. This supports the finite-sample stability of the empirical distance matrix, although it does not constitute a proof of QAP-permutation consistency.

### 5.4 Study 3: permutation test for label-dependent topology

#### Design

Study 3 tests whether the S-IGTD topology depends on the true class labels rather than on label-independent structure or QAP randomness. We used *p* = 400 with a 20 × 20 grid, 36 signal features arranged as three blocks of 12, *n*_train_ = 600 with 100 samples per class, *n*_test_ = 600, and *ρ*_noise_ = 0.80. Nuisance features used nominal block size 10, giving 36 nuisance blocks, four with 11 features and 32 with 10 features, nuisance scale *ν* = 1.0, and no additional latent factors. The observed statistic was

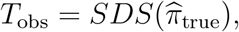

where 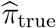 is the S-IGTD permutation learned from the true labels. For the null distribution, we independently permuted the training labels and re-ran the full S-IGTD pipeline, including topology optimization, for *B* = 500 null replicates. Since lower SDS is better, the empirical one-sided permutation *p*-value is

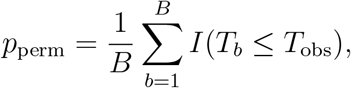

where *T*_*b*_ is the SDS from null replicate *b*.

#### Result interpretation

Figure 4 shows that the observed SDS, *T*_obs_ = 4.951, lies in the left tail of the label-randomized null distribution, whose mean is 7.679. Because lower SDS is better, the left-tail position indicates stronger signal-feature compactness than expected when topology construction is label-randomized. The empirical permutation value is *p*_perm_ = 0.032, so the test rejects the label-randomized null at the 5% level. This supports the conclusion that S-IGTD learns label-dependent, class-informative topology. This permutation analysis uses one simulated dataset with 500 label permutations; it is a label-dependence test for topology construction, not a multi-split classifier benchmark.

**Figure 4.**
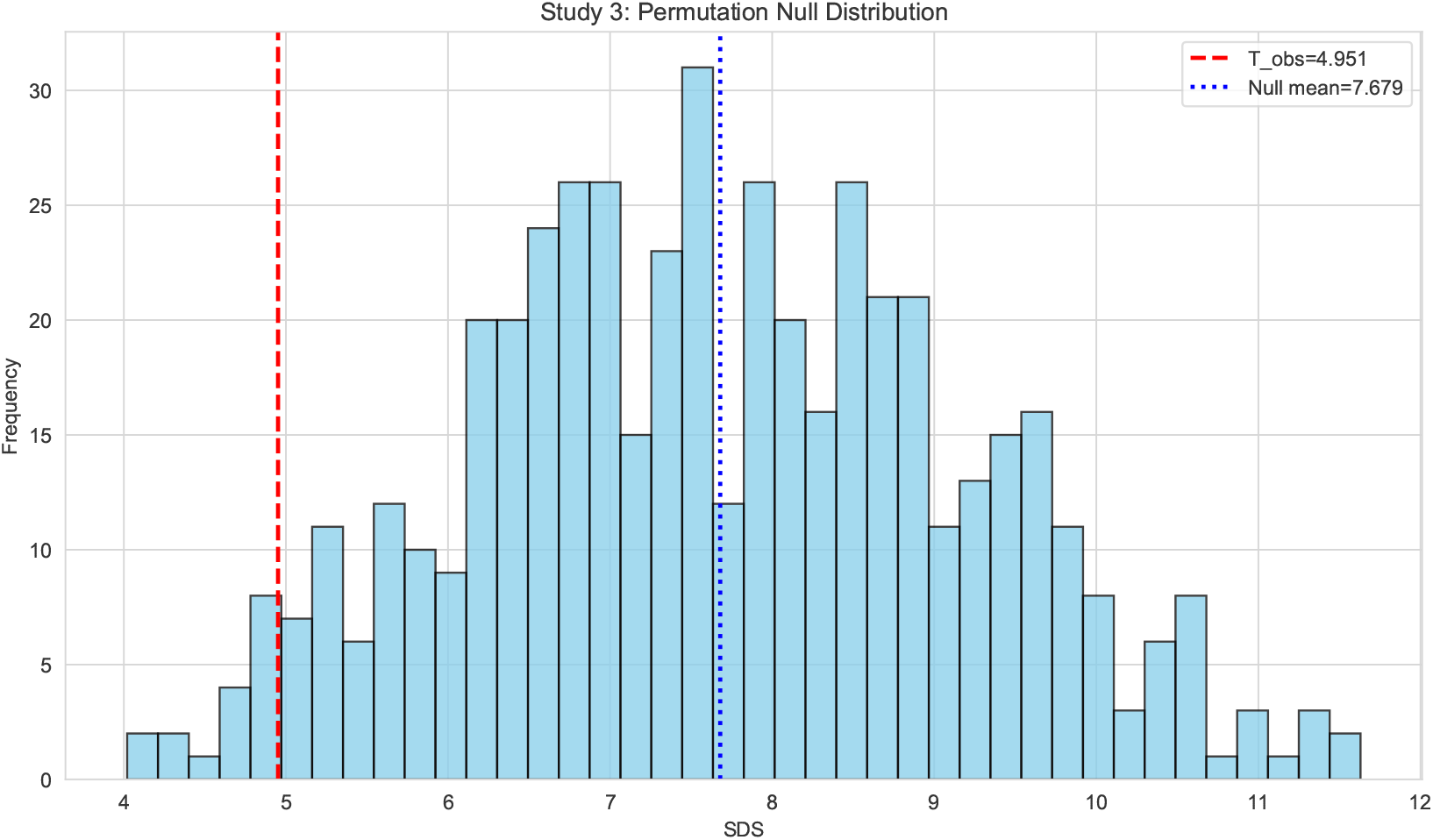
Study 3 permutation-null distribution of SDS under label randomization. The vertical line marks the observed SDS obtained from the true labels. Because lower SDS is better, an observed value in the left tail indicates stronger signal-feature compactness than expected under label-randomized topology construction.

### 5.5 Study 4: joint scaling of dimensionality and signal mass

#### Design

Study 4 evaluates whether the topology advantage persists as the feature dimension increases. We fixed the training size at *n*_*k*_ = 10 per class, so *n*_train_ = 60, and used *n*_test_ = 600.

The feature dimension was varied as

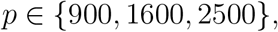

with grids 30×30, 40×40 and 50×50, respectively. The signal fraction was fixed at 12% and split equally across the three signal blocks, giving 108, 192 and 300 signal features. Nuisance features used nominal block size 200, nuisance scale *ν* = 4.0, six class-independent latent factors with latent scale 2.5, and *ρ*_noise_ = 0. This gives three nuisance blocks of 264 features at *p* = 900, seven blocks at *p* = 1600 (one of size 202 and six of size 201), and eleven blocks of 200 features at *p* = 2500. Each setting used 10 Monte Carlo replications. Because the signal fraction is held constant as *p* grows, the absolute number of signal features and the total signal mass both increase with *p*. Therefore, this study jointly scales feature dimension and signal mass rather than isolating a pure high-dimensional-low-sample-size setting.

#### Result interpretation

Figure 5 shows that S-IGTD has the lowest mean SDS at all tested dimensions. The SDS advantage is Holm-adjusted significant for all three *p* values. S-IGTD also has higher mean accuracy and macro-F1 score, but the classifier comparisons at *p* = 1600 are directional rather than Holm-adjusted significant under the paired tests. We therefore interpret Study 4 primarily as a topology result because the design jointly increases dimension and signal mass: the between-class distance continues to produce more compact signal layouts as the grid size grows.

**Figure 5.**
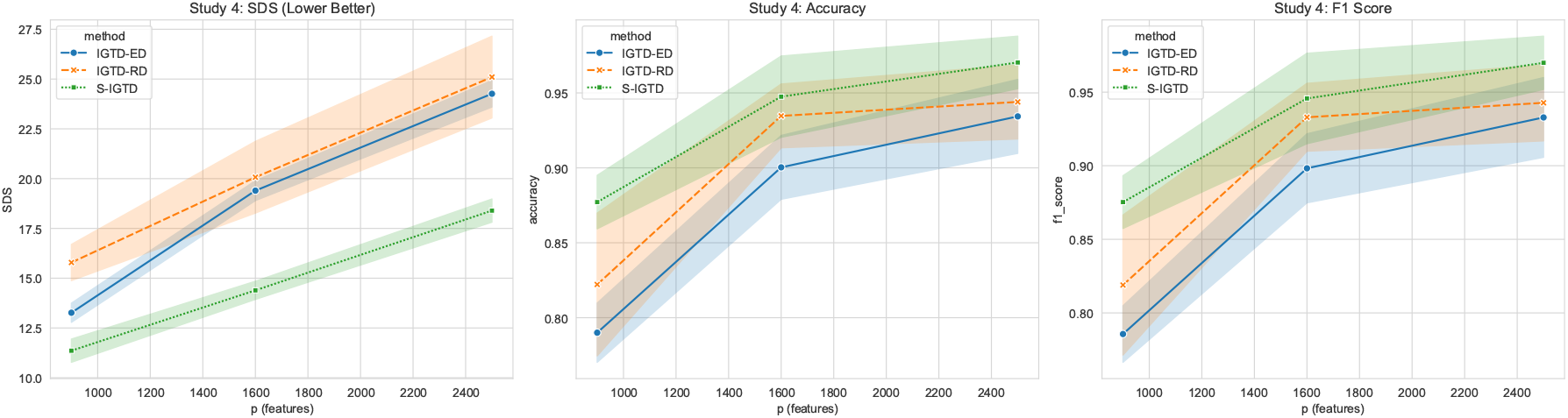
Study 4 trends under joint scaling of feature dimension and signal mass. Lines show Monte Carlo means across replications; shaded bands are the seaborn default 95% confidence intervals around the mean. Numerical summaries are reported in Table 2.

Overall, the simulations support the topology-focused claim. Constructing the image layout from between-class correlation yields more compact signal-feature neighborhoods under strong nuisance correlation, finite training samples and increasing feature dimension. Downstream classifier performance also improves in mean in these controlled settings, but accuracy and macro-F1 score remain secondary diagnostics rather than claims of general classifier superiority. The broader benchmark against projection-based tabular-to-image methods is reported in Section 6.

## 6 Real data analysis

We conducted the real-data benchmark on five public biological datasets. The primary outcome is topology compactness measured by SDS. Accuracy and the macro-F1 score are reported as downstream classifier diagnostics, because Sections 4–5 support the topology objective rather than classifier optimality. A single Conv-BN-ReLU classifier was used for all real-data image layouts so that differences in downstream performance reflect the constructed topology rather than architecture selection. For fairness, optimization limits and sampling caps were held fixed within each family of image-layout methods.

### 6.1 Datasets and preprocessing

The benchmark covers five data modalities, ordered in Table 3 as they are introduced here: bulk RNA-seq pan-cancer profiling (TCGA), single-cell RNA-seq (PBMC 3k), microbiome relative-abundance profiles (HMP Body Sites), DNA methylation (MethPed), and metabolomics (MultiClassMetabo). This order groups the datasets by biological assay rather than alphabetically or by sample size. The main manuscript reports the post-filter analysis dimensions; dataset-specific preprocessing details are provided in Supplementary Note S1.

**Table 3.** Dataset sources and metadata after filtering and feature selection.

| Dataset | Source | Samples ($n$) | Features ($p$) | Classes ($K$) | Grid |
| --- | --- | --- | --- | --- | --- |
| TCGA Pan-Cancer | TCGA Pan-Cancer Atlas / UCSC Xena (Weinstein et al., 2013; Hoadley et al., 2018; Goldman et al., 2020) | 11,014 | 2,500 | 33 | $50 \times 50$ |
| PBMC 3k | 10x Genomics via Scanpy (10x Genomics, 2016; Wolf et al., 2018) | 2,699 | 2,500 | 7 | $50 \times 50$ |
| HMP Body Sites | HMP via `curatedMetagenomicData` (The Human Microbiome Project Consortium, 2012; Pasolli et al., 2017) | 748 | 740 | 5 | $20 \times 37$ |
| MethPed | GEO GSE90496 CNS methylation reference cohort (Capper et al., 2018; NCBI Gene Expression Omnibus, 2018) | 2,801 | 2,500 | 91 | $50 \times 50$ |
| MultiClassMetabo | MetaboLights MTBLS52 (Haug et al., 2020; MetaboLights, 2012) | 32 | 104 | 4 | $8 \times 13$ |

All datasets were processed by deterministic loaders before stratified 80*/*20 training–test partitioning. Briefly, TCGA and MethPed used variance-based feature retention to fit the 50 × 50 image grid, PBMC 3k used standard single-cell normalization and highly variable gene selection, and HMP Body Sites and MultiClassMetabo were used without truncation because their feature counts were already below the grid cap. Min–max scaling to [0, 1] was fit on each training set and applied to the corresponding test set.

### 6.2 Experimental protocol

The tabular-to-image literature is broad, and no single layout family is expected to provide the best result on every dataset (Shwartz-Ziv and Armon, 2022). We therefore compared S-IGTD with representative layout families rather than attempting an exhaustive leader-board. Random and Identity provide sanity controls; IGTD-ED and IGTD-RD represent the optimization-based IGTD family; TINTO-DeepInsight-PCA, TINTO-DeepInsight-tSNE and TINTO-REFINED-MDS represent projection- or MDS-based generators implemented through a common reproducible interface (Castillo-Cara et al., 2023; Liu et al., 2025). Each dataset used 10 stratified 80*/*20 training–test replications with matched random seeds across methods.

We also include F-stat-RowMajor as a supervised univariate reference for SDS. For each training set, let *F*_*j*_ denote the one-way ANOVA statistic for feature column **X**_·*j*_ against the class labels. F-stat-RowMajor sorts features by decreasing *F*_*j*_ and places them on the image grid in row-major order. Because the real-data SDS proxy is defined from the same top-ranked ANOVA features, this reference is designed to give a favorable univariate layout for SDS. It is therefore not treated as a fair competitor to S-IGTD’s pairwise topology construction, but as a calibration point for interpreting the SDS scale.

The S-IGTD estimator in Section 4 applies to multiclass settings. It was used for TCGA Pan-Cancer, PBMC 3k, HMP Body Sites, MethPed and MultiClassMetabo. The same optimization limits and sample caps were used for competing image-layout methods within each method family; exact computational settings are listed in Supplementary Note S2.

The downstream classifier for the main benchmark was a three-stage Conv→BatchNorm→ReLU stack with adaptive average pooling and a linear head, trained from scratch in PyTorch for 50 epochs (batch size 32). This classifier was selected because a feature-layout method should be evaluated with a CNN that can use local spatial structure.

Only one CNN architecture was used in the main real-data benchmark. This is intentional: the experiment evaluates whether different feature-to-pixel layouts provide usable local structure for a fixed downstream CNN, not whether one classifier architecture dominates another. A broader classifier benchmark, including LeNet-5 and non-CNN tabular learners, would address a different question and is left to future comparative work.

Because true signal indices are not available for real biological data, we compute SDS against a label-derived proxy: the top-12% of features by training-set ANOVA *F* -statistic, with a lower bound of 20 and an upper bound of 300 features. The 12% rule was pre-specified to retain a sufficiently large signal proxy for stable pairwise-distance measurement while avoiding a proxy set so broad that it becomes insensitive to local compactness. Real-data SDS must therefore be interpreted as *agreement with a supervised univariate proxy*, not as a ground-truth measure of biological signal recovery. Because grid dimensions differ substantially across datasets (e.g. 50 × 50 for TCGA versus 8 × 13 for MultiClassMetabo), we avoid cross-dataset averaging of raw SDS in Table 4 and instead report per-dataset Holm-adjusted significant win counts. Accuracy and the macro-F1 score are reported as downstream classifier diagnostics. Supplementary Note S3 gives the validation metric definitions.

**Table 4.** Dataset-level real-data benchmark summary. Lower SDS and negative Δ favor S-IGTD.

| Dataset | $K$ | Wins | SDS | $\Delta_{RD}$ | Acc. |
| --- | --- | --- | --- | --- | --- |
| TCGA | 33 | 7/7 | 22.97 | -3.48 | 0.932 |
| PBMC | 7 | 6/7 | 25.67 | -1.77 | 0.895 |
| MethPed | 91 | 6/7 | 24.83 | -0.79 | 0.951 |
| HMP | 5 | 4/7 | 13.30 | -0.95 | 0.922 |
| Metabo | 4 | 1/7 | 4.94 | -0.18 | 0.400 |

Wins are Holm-adjusted significant SDS reductions versus seven non-reference controls. $\Delta_{RD}$ is the median paired SDS difference, S-IGTD minus IGTD-RD. Acc. is mean Conv-BN-ReLU accuracy.

Statistical testing used paired Wilcoxon signed-rank tests over the 10 matched replications (S-IGTD vs. each comparator, per dataset and per metric), with one-sided alternatives pre-specified: “less” for SDS and “greater” for accuracy and the macro-F1 score. Holm-Bonferroni correction (Holm, 1979) was applied within each dataset and metric family at *α* = 0.05. We use “Holm-adjusted significant” to mean Holm-adjusted *p <* 0.05. Two-sided sensitivity analyses and per-comparison effect-size estimates with 95% bootstrap confidence intervals are included in the released result tables.

### 6.3 Topology compactness: headline SDS results

The core benchmark result is class-supervised topology compactness. Excluding the supervised F-stat-RowMajor reference, S-IGTD achieved 24*/*35 Holm-adjusted significant SDS wins against seven non-reference layout controls (5 datasets × 7 controls). The per-dataset breakdown is: TCGA Pan-Cancer 7*/*7, PBMC 3k 6*/*7, MethPed 6*/*7, HMP Body Sites 4*/*7, and MultiClassMetabo 1*/*7. Restricted to the four larger multiclass benchmarks (TCGA, PBMC, MethPed and HMP), the count is 23*/*28; the very-small-sample MultiClassMetabo dataset is reported as a sensitivity benchmark. Median SDS reductions of S-IGTD versus IGTD-RD are largest on TCGA (Δ = −3.48, Holm-adjusted *p* = 0.047) and PBMC (Δ = −1.77, *p* = 0.047) and smaller but significant on HMP (Δ = −0.95, *p* = 0.049) and MethPed (Δ = −0.79, *p* = 0.047).

The comparison separates three kinds of controls. Against simple and IGTD-family controls, S-IGTD has lower mean SDS than Random and IGTD-ED on all five datasets and lower mean SDS than IGTD-RD on four of five datasets. Against projection-based TINTO methods, S-IGTD has lower mean SDS than TINTO-DeepInsight-PCA on four of five datasets, TINTO-DeepInsight-tSNE on three of five, and TINTO-REFINED-MDS on five of five. The largest S-IGTD reductions occur on TCGA Pan-Cancer and MethPed, the large multiclass datasets where class-centroid correlation is most informative.

The F-stat-RowMajor reference attains the lowest absolute SDS on every dataset by construction because it packs the top-ANOVA features into adjacent pixels in row-major order. S-IGTD is not intended to beat this univariate reference on its own proxy; rather, F-stat-RowMajor provides a calibration point for how low the proxy SDS can be when pairwise topology is ignored. Table 4 summarizes the dataset-level results.

A centroid-weighting sensitivity analysis compared the class-balanced centroid estimator used by S-IGTD with a class-probability-weighted variant. On real, generally unbalanced datasets, the class-balanced version attained Holm-adjusted significantly lower SDS in all five datasets at *α* = 0.05 (TCGA Δ = −3.58, *p*_Holm_ = 0.012; PBMC Δ = −1.23, *p*_Holm_ = 0.012; MethPed Δ = −2.09, *p*_Holm_ = 0.012; HMP Δ = −0.73, *p*_Holm_ = 0.027; MultiClassMetabo Δ = −0.63, *p*_Holm_ = 0.012). This sensitivity analysis supports the estimator choice in Section 3.2: S-IGTD uses a balanced surrogate for the between-group correlation target, and the guarantees in Section 4 are stated for that surrogate. Table 5 reports the per-dataset SDS for both centroid variants alongside IGTD-RD as an unsupervised reference.

**Table 5.**
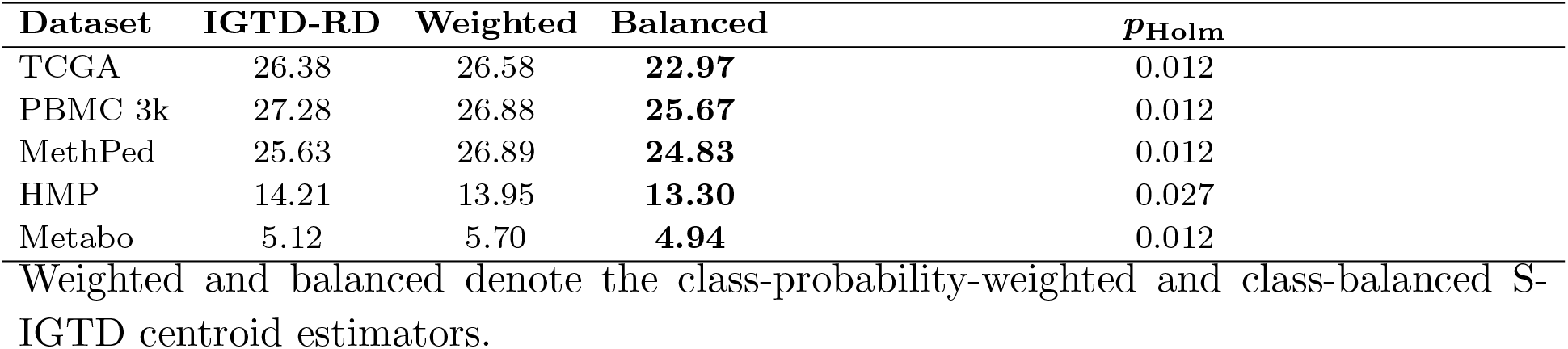
Centroid-weighting sensitivity analysis. Mean SDS over 10 stratified replications; lower is better. *p*_Holm_ compares balanced versus weighted S-IGTD by paired Wilcoxon test.

### 6.4 Classifier compatibility under Conv-BN-ReLU

Topology compactness is the primary outcome, but a useful tabular-to-image representation should also be usable by a CNN. Under Conv-BN-ReLU, S-IGTD has higher mean accuracy than Random and IGTD-RD on all five datasets and higher mean accuracy than IGTD-ED on four of five datasets. These are mean comparisons. After Holm correction over matched replications, however, the accuracy tests do not support a uniform claim that S-IGTD is significantly more accurate than all IGTD-style layouts. This distinction matters because the real-data benchmark has only 10 matched replications, and one dataset is a sensitivity case with a very small test set. Relative to projection-based TINTO methods, S-IGTD has higher mean accuracy than TINTO-DeepInsight-PCA on four of five datasets, TINTO-DeepInsight-tSNE on two of five datasets, and TINTO-REFINED-MDS on four of five datasets. Table 6 reports the full per-dataset accuracy panel.

**Table 6.** Mean classification accuracy under the Conv-BN-ReLU classifier (50 epochs, 10 stratified replications). F-stat-RowMajor is a supervised univariate reference that places features in descending training-set ANOVA-*F* order. **Bold** is the per-dataset best mean accuracy; underlined marks S-IGTD when it is not best. Accuracy is treated as a downstream CNN diagnostic rather than as the primary benchmark metric.

| Dataset | F-stat | Random | Identity | IGTD-ED | IGTD-RD | S-IGTD | TINTO-PCA | TINTO-tSNE | TINTO-MDS |
| --- | --- | --- | --- | --- | --- | --- | --- | --- | --- |
| TCGA ($K=33$) | 0.932 | 0.932 | **0.933** | 0.931 | 0.932 | <u>0.932</u> | 0.930 | 0.932 | 0.930 |
| PBMC 3k ($K=7$) | **0.924** | 0.889 | 0.893 | 0.880 | 0.893 | <u>0.895</u> | 0.869 | 0.899 | 0.869 |
| HMP ($K=5$) | 0.908 | 0.897 | 0.908 | 0.917 | 0.908 | <u>0.922</u> | **0.928** | 0.923 | 0.903 |
| MethPed ($K=91$) | **0.951** | 0.947 | 0.947 | 0.949 | 0.949 | **0.951** | 0.938 | 0.937 | 0.938 |
| MultiClassMetabo ($K=4$) | **0.414** | 0.314 | 0.343 | 0.386 | 0.357 | <u>0.400</u> | 0.329 | **0.414** | 0.357 |

On MethPed, the highest-class-count benchmark (*K* = 91), S-IGTD matches the supervised univariate F-stat-RowMajor reference in mean accuracy (0.951 versus 0.951; paired Wilcoxon *p* = 0.628 for the paired difference). This is a favorable compatibility result in the setting where class-centroid covariation is expected to be most informative, but it remains a secondary CNN diagnostic rather than a primary topology claim. On TCGA Pan-Cancer, S-IGTD also reaches near-best accuracy (0.932 versus 0.933 for Identity). MultiClassMetabo has only *n* = 32 samples, so the per-replication test set is very small; its accuracy differences are reported as a small-sample sensitivity result rather than a quantitative ranking.

The accuracy panel should be interpreted as dataset-dependent. S-IGTD is a lower-SDS alternative to IGTD-style layouts and has comparable mean accuracy to projection-based TINTO layouts, but it is not uniformly superior to every projection method. TINTO-DeepInsight-tSNE remains an important comparator, especially on smaller or low-class-count datasets, where global projection geometry can be effective.

Overall, the real-data benchmark supports a topology-focused contribution: S-IGTD produces compact supervised layouts relative to IGTD-style and other non-reference controls, while F-stat-RowMajor remains a supervised univariate SDS reference. Under Conv-BN-ReLU, S-IGTD has higher mean accuracy than IGTD-style layouts on most datasets, but the corrected accuracy tests do not support a uniform accuracy-superiority claim. This pattern is consistent with the design of the method: S-IGTD is best supported as a class-supervised topology generator, especially for large multiclass biological datasets, while projection-based TINTO methods remain important comparators on smaller or low-class-count datasets.

## 7 Conclusion and discussion

S-IGTD replaces IGTD’s unsupervised feature-distance metric with the supervised distance *d*_sup_ = 1 −|*ρ*_*B*_| on the class-centroid matrix, motivated by the Within-And-Between-Analysis decomposition. The contribution is to integrate class supervision at the pairwise distance level rather than as univariate feature pre-processing, addressing an open methodological gap in tabular-to-image methods.

Theoretically, we establish entrywise consistency of the supervised distance matrix and identify balanced-class signal/noise settings in which the supervised metric improves the SDS-related topology objective. Empirically, S-IGTD produced more compact supervised layouts than IGTD-style and other non-reference controls across the biological benchmarks, with the strongest evidence on large multiclass datasets where class-centroid covariation is most informative. Accuracy results are secondary: under a Conv-BN-ReLU CNN, S-IGTD has higher mean accuracy than IGTD-style baselines on most datasets, but the corrected accuracy tests do not support a uniform accuracy-superiority claim. This dataset-dependent interpretation is consistent with the broader tabular-to-image literature, where projection-based, optimization-based and toolkit implementations make different layout assumptions and no single family is expected to dominate all datasets (Jiang et al., 2025; Shwartz-Ziv and Armon, 2022).

Four limitations warrant explicit discussion. First, the downstream Quadratic Assignment Problem is NP-hard and is the dominant runtime cost of both IGTD and S-IGTD; the S-IGTD distance step itself is *O*(*Kp*^2^) and does not add overhead beyond IGTD. Second, Pearson correlation across class centroids captures linear class-mean covariation rather than arbitrary nonlinear class structure. Third, the fixed rectangular grid imposes geometric constraints and can require padding when *p* does not factorise conveniently. Fourth, real-data SDS uses an ANOVA-*F* proxy because ground-truth signal features are unknown, so SDS measures agreement with a supervised univariate topology proxy rather than direct biological signal recovery.

S-IGTD is most useful when a class-supervised image layout is the analysis target, especially in multiclass omics settings with informative class-centroid covariation. The resulting pseudo-images can be analyzed by CNNs or by other downstream methods that use spatial neighborhood structure, image texture or local feature aggregation. For very high-dimensional tables, feature screening before QAP layout may still be needed. When global feature geometry is the dominant structure of interest, projection-based methods such as DeepInsight or REFINED may be equally appropriate or preferable.

Three directions follow naturally. *Dynamic S-IGTD* would treat the permutation matrix as a learnable parameter via differentiable ranking, allowing topology and classifier to co-train. *Regression S-IGTD* would replace the class-centroid matrix with conditional expectations over quantile strata for continuous outcomes. *Classifier-aware topology* would optimize arrangement for a target architecture’s local-filter statistics. Additional methodological work should evaluate signed rather than absolute between-group correlation and residualized WABA estimators that more directly remove within-class covariance.

## Data availability

The Python source code, dataset preprocessing scripts, analysis configuration files, seed manifests, per-replication result tables and figure-generation scripts used in this paper are available from the GitHub repository at https://github.com/hanmingwu1103/S-IGTD. The public benchmark datasets are cited in Table 3 and in the reference list; processed analysis-ready files and reproducibility instructions are provided with the repository for peer review and reuse.

