## Supplementary Material for "S-IGTD: supervised tabular-to-image topology learning via between-group correlation for multiclass classification of biological data"

May 18, 2026

#### Supplementary Note S1: Dataset Sources

Table 1 lists the public data sources used in the real-data benchmark. The post-filter dimensions match Table 3 of the main manuscript. When an analysis-ready CSV was used by the benchmark script, the CSV was generated from the listed source by the released dataset-building scripts.

#### Supplementary Note S2: Dataset Preprocessing

All real-data benchmarks used deterministic preprocessing before the stratified training–test partitions described in the main manuscript. Non-finite feature values were replaced during loading, features with no usable variation were removed or assigned zero contribution by the downstream scaling rule, and class labels were encoded after dataset-specific cleaning. Min–max scaling to $[0, 1]$ was fit on each training set and then applied to the corresponding test set, so no test-set information entered the scaling parameters.

*

Table 1: Public dataset sources and analysis dimensions.

| Dataset | Modality | Source link | $n$ | $p$ | $K$ | Grid |
|---|---|---|---|---|---|---|
| TCGA Pan-Cancer | Bulk RNA-seq | UCSC Xena Pan-Cancer Atlas download hub: `https://pancanatlas.xenahubs.net/download/` | 11,014 | 2,500 | 33 | $50 \times 50$ |
| PBMC 3k | Single-cell RNA-seq | 10x Genomics 3k PBMC dataset: `https://www.10xgenomics.com/datasets/3-k-pbm-cs-from-a-healthy-donor-1-standard-1-1-0`; accessed through Scanpy's `pbmc3k` loader | 2,699 | 2,500 | 7 | $50 \times 50$ |
| HMP Body Sites | Microbiome relative abundance | Human Microbiome Project profiles through Bioconductor `curatedMetagenomicData`: `https://bioconductor.org/packages/curatedMetagenomicData` | 748 | 740 | 5 | $20 \times 37$ |
| MethPed | DNA methylation | GEO accession GSE90496: `https://www.ncbi.nlm.nih.gov/geo/query/acc.cgi?acc=GSE90496`; beta matrix: `https://ftp.ncbi.nlm.nih.gov/geo/series/GSE90nnn/GSE90496/suppl/GSE90496_beta.txt.gz` | 2,801 | 2,500 | 91 | $50 \times 50$ |
| MultiClassMetabo | Metabolomics | MetaboLights MTBLS52: `https://www.ebi.ac.uk/metabolights/MTBLS52`; public study files under `https://ftp.ebi.ac.uk/pub/databases/metabolights/studies/public/MTBLS52/` | 32 | 104 | 4 | $8 \times 13$ |

**TCGA Pan-Cancer.** The TCGA Pan-Cancer RNA-seq matrix was downloaded from the UCSC Xena Pan-Cancer Atlas hub using the expression file `EB++AdjustPANCAN_IlluminaHiSeq_RNASeqV2.geneExp.xena.gz`. Cancer-type labels were obtained from the Xena phenotype file `Survival_SupplementalTable_S1_20171025_xena_sp.gz`. Samples were aligned between expression and phenotype tables, and the analysis retained the 2,500 genes with largest empirical variance. The resulting analysis matrix had $n = 11{,}014$ samples, $p = 2{,}500$ features, $K = 33$ cancer-type classes and a $50 \times 50$ grid.

**PBMC 3k.** The PBMC 3k single-cell RNA-seq dataset was loaded through the standard Scanpy dataset interface. The preprocessing pipeline filtered low-information cells and genes, applied total-count normalization to a target sum of 10,000, applied the `log1p` transformation, and selected 2,500 highly variable genes. The released loader then used deterministic clustering labels for the benchmark label column. After rare-cluster filtering, the resulting analysis matrix had $n = 2{,}699$ cells, $p = 2{,}500$ genes, $K = 7$ classes and a $50 \times 50$ grid.

**HMP Body Sites.** The HMP Body Sites microbiome profile was constructed from the Human Microbiome Project relative-abundance data available through `curatedMetagenomicData`. The loader encoded body-site labels, retained five body-site classes, and used the available relative-abundance features after deterministic cleaning. The feature count was already below the 2,500-feature cap, so no top-variance truncation was applied. The resulting analysis matrix had $n = 748$ samples, $p = 740$ features, $K = 5$ body-site classes and a $20 \times 37$ grid.

**MethPed.** The MethPed methylation benchmark was obtained from GEO accession GSE90496. The loader used the processed supplementary beta-value matrix, aligned sample columns with the GEO phenotype table, extracted the methylation-class label, and retained the 2,500 CpG probes with largest empirical variance from the processed beta matrix used for the benchmark. The resulting analysis matrix had $n = 2{,}801$ samples, $p = 2{,}500$ features, $K = 91$ classes and a $50 \times 50$ grid.

**MultiClassMetabo.** The MultiClassMetabo benchmark was derived from MetaboLights study MTBLS52. The loader read the metabolite assignment file and sample metadata, matched abundance columns to sample identifiers, used the available disease-stage or related study label column from the sample metadata, and replaced missing abundance values by a conservative feature-table value before analysis. The feature count was already below the feature cap, so no top-variance truncation was applied in the benchmark. The resulting analysis matrix had $n = 32$ samples, $p = 104$ features, $K = 4$ classes and an $8 \times 13$ grid. Because the test set is very small in each stratified partition, classifier results for this dataset are interpreted as a sensitivity analysis.

### Supplementary Note S3: Benchmark Protocol

Each dataset used 10 stratified 80/20 training–test replications with matched random seeds across feature-layout methods. Within a replication, all methods used the same training set, test set, scaling parameters and class-label encoding. The benchmark compared S-IGTD with Random, Identity, IGTD-ED, IGTD-RD, F-stat-RowMajor, TINTO-DeepInsight-PCA, TINTO-DeepInsight-tSNE and TINTO-REFINED-MDS.

Random used a uniform-random feature permutation. Identity used the original feature order and placed features row-major. IGTD-ED and IGTD-RD used the IGTD optimization framework with Euclidean and total-correlation feature distances, respectively. S-IGTD used the class-balanced centroid-correlation distance for multiclass datasets. F-stat-RowMajor sorted features by decreasing training-set ANOVA $F$-statistic and then placed features row-major. The TINTO comparators represented projection- or MDS-based tabular-to-image generators.

For S-IGTD and the IGTD-style comparators, the maximum number of QAP optimization iterations was fixed at 300 in the real-data benchmark. For topology-computing methods that subsample samples when estimating pairwise feature structure, the maximum number of samples used for topology construction was fixed at 1,200. These limits were held fixed across competing image-layout methods so that the comparison reflected the feature-distance definition rather than different optimization effort.

The downstream classifier in the real-data benchmark was a three-stage Conv-BN-ReLU CNN with adaptive average pooling and a linear head, trained from scratch for 50 epochs with batch size 32. This classifier was used as a fixed downstream diagnostic for image-layout compatibility; accuracy and the macro-F1 score were not treated as the primary endpoint.

### Supplementary Note S4: Real-Data SDS Proxy and Validation Metrics

On simulated data, SDS is computed using the known signal-feature index set $\mathcal{S}_{\text{sig}}$. On real biological datasets, ground-truth signal indices are unavailable. The main manuscript therefore uses a supervised univariate proxy set

$$\mathcal{S}_F = \{j : \; F_j \text{ is among the selected largest training-set ANOVA statistics}\},$$

where $F_j$ is the one-way ANOVA statistic for feature column $\mathbf{X}_{\cdot j}$ against the training labels. The number of selected features is

$$s = \max\{20, \min(300, \lfloor 0.12p \rfloor)\}.$$

The lower bound avoids an unstable SDS estimate based on too few features, and the upper bound avoids a proxy set that is so large that it becomes insensitive to local compactness. The resulting real-data SDS measures agreement with a supervised univariate feature proxy, not recovery of known biological ground-truth signal features.

For a learned feature-to-pixel permutation $\widehat{\pi}$, the real-data SDS proxy is

$$\mathrm{SDS}_F(\widehat{\pi}) = \frac{2}{s(s-1)} \sum_{u \in \mathcal{S}_F} \sum_{\substack{v \in \mathcal{S}_F \\ v > u}} \left\| \mathrm{loc}_{\widehat{\pi}}(u) - \mathrm{loc}_{\widehat{\pi}}(v) \right\|_2 .$$

Lower values indicate that the selected supervised proxy features are more spatially compact.

Classification accuracy is the fraction of correctly classified test-set samples. The macro-F1 score is the unweighted mean of class-specific F1 scores. Accuracy and the macro-F1 score are downstream CNN diagnostics in the real-data benchmark; they are not the primary endpoint of the topology method.

#### Supplementary Note S5: Statistical Testing

For each dataset and metric, S-IGTD was compared with each comparator over the same 10 stratified training–test replications. Paired Wilcoxon signed-rank tests used one-sided alternatives: "less" for SDS and "greater" for accuracy and the macro-F1 score. Holm-Bonferroni correction was applied within each dataset and metric family. A result is described as Holm-adjusted significant when the adjusted $p$-value is below 0.05. The released result tables also include two-sided sensitivity analyses and bootstrap confidence intervals for paired effect-size summaries.

#### Supplementary Note S6: Reproducibility Files

The public repository contains the source code, dataset-building scripts, analysis configuration files, seed manifests, per-replication result tables and figure-generation scripts. The main reproducibility entry points are the dataset builders in `Real_Data_Analysis/`, the real-data benchmark script `real_data_studies.py`, the simulation scripts and locked result tables, and the manuscript figure-generation scripts. Processed analysis-ready CSV files are provided with the repository for peer review and reuse when direct database access is inconvenient.
